# The emerging British *Verticillium longisporum* population consists of aggressive *Brassica* pathogens

**DOI:** 10.1101/111922

**Authors:** Jasper R. L. Depotter, Luis Rodriguez-Moreno, Bart P.H.J. Thomma, Thomas A. Wood

**Affiliations:** Laboratory of Phytopathology, Wageningen University and Research, Droevendaalsesteeg 1, 6708 PB Wageningen, The Netherlands; Department of Crops and Agronomy, National Institute of Agricultural Botany, Huntingdon Road, CB3 0LE Cambridge, United Kingdom

**Keywords:** Verticillium stem striping, Verticillium wilt, oilseed rape, pathogenicity, hybrid, cauliflower, cabbage, *Arabidopsis*

## Abstract

The impact of diseases depends on the dynamic interplay between host, pathogen and the environment. Newly emerging diseases may be the consequence of novel pathogen introductions that are typically associated with unpredictable outcomes, as their interaction with the host in a novel environment is unprecedented. Alternatively, new diseases may emerge from latent, previously established, pathogen populations that are triggered by changes in environmental factors like weather, agricultural practices and ecosystem management. Verticillium stem striping recently emerged in British oilseed rape (*Brassica napus*) production from a latent *Verticillium longisporum* population. *V. longisporum* is a hybrid fungal pathogen consisting of three lineages, each representing a separate hybridization event. Despite its prevalence, little is known of the pathogenicity of the British *V. longisporum* population. In this study, the pathogenicity of British isolates was tested on four different cultivars of three different *Brassica* crop species as well as on the model plant *Arabidopsis thaliana* and compared with previously characterized *V. longisporum* strains from other regions of the world, including representatives of all three hybrid lineages. Intriguingly, the British isolates appeared to be amongst the most pathogenic strains on *Brassica* crops. In conclusion, Verticillium stem striping poses a genuine threat to oilseed rape production as the British *V. longisporum* population consists of aggressive pathogens that have the potential to significantly impact *Brassica* crops.

## Introduction

*Verticillium* fungi cause wilt diseases on hundreds of plant species of which many are economically important crops. *Verticillium dahliae* is the most notorious member of these fungi and can cause severe yield losses in crops like olive and cotton (Fradin and Thomma 2006; Levin et al. 2003; Melero-Vara et al. 1995). Despite a wide host range that comprises hundreds of plant species, *V. dahliae* generally does not infect brassicaceous plants (Depotter *et al.,* 2016a). In contrast, *V. longisporum* is specialized on Brassicaceae hosts, with oilseed rape as its most economically important target (Depotter *et al.,* 2016a; Inderbitzin *et al.,* 2011). In contrast to all other *Verticillium* spp, including *V. dahliae, V. longisporum* is not a haploid organism but rather an allopolyploid as a consequence of interspecific hybridization. The species *V. longisporum* consists of three lineages, each representing a separate hybridization event. Four parental lines, including two *V. dahliae* isolates (D2 and D3), contributed to the different hybridization events. The two remaining parental lines represent two previously uncharacterized *Verticillium* species that have been provisionally called Species A1 and Species D1 (Inderbitzin et al. 2011). Species A1 participated in all three hybridization events and hybridized with D1, D2 and D3 to form the lineages A1/D1, A1/D2 and A1/D3, respectively. Conceivably, the allopolyploid genome of *V. longisporum* contributed to the host range shift such that *V. longisporum* gained the capacity to infect Brassicaceae (Inderbitzin et al. 2011; Depotter et al. 2016b).

Interspecific hybridization is an important driver for genome evolution (Depotter *et al.,* 2016b). Hybrid organisms experience a so-called “genomic shock” that incites major genomic rearranges and changes in gene expression patterns (Doyle et al. 2008). Hybrids are therefore especially adept in responding to environmental changes or invading novel niches. Hybridization brings important additions to the toolset of pathogens; hence, host range and pathogenicity are often altered in interspecific hybrids (Depotter et al. 2016b). Differences between *V. longisporum* lineages may therefore originate from their separate hybrid origin. Lineages A1/D1 and A1/D3 are found on various Brassicaceae species, whereas lineage A1/D2 is only known from horseradish in the USA (Inderbitzin et al. 2011; Yu et al. 2016). A1/D1 is the predominant lineage on oilseed rape and is also the most pathogenic *V. longisporum* lineage of this crop (Novakazi et al. 2015).

In addition to its different genetic constitution, *V. longisporum* is unique amongst Verticillium species for its disease symptom display on oilseed rape. Verticillium pathogens are xylem colonizers inducing occlusions in the vessels, which hampers the water transport in the xylem (Fradin and Thomma 2006). In response, Verticillium infections generally develop wilting symptoms. However, these symptoms are lacking from *V. longisporum* infections on oilseed rape. Rather, black unilateral stripes appear on the plant stem at the end of the growing season and, in a later stage, microsclerotia appear on the cortex under the stem epidermis (Heale and Karapapa 1999; Depotter et al. 2016a). Hence, the new common name “Verticillium stem striping” was coined to describe the *V. longisporum* disease on oilseed rape (Depotter et al. 2016a). Intriguingly, Verticillium stem striping symptoms fail to appear during pathogenicity tests when oilseed rape plants are grown under controlled conditions and seedlings are inoculated by dipping the roots in a spore suspension (Eynck et al. 2007; Eynck et al. 2009; Floerl et al. 2008; Zeise and Tiedemann 2002). Under those conditions, plants exhibit chlorosis, vascular discoloration and stunting at an early stage. The reasons for these differences in symptom development are currently unknown. Nevertheless, despite differences in disease symptomatology, it has previously been determined that root-dip pathogenicity tests in the glasshouse are a good proxy for oilseed rape cultivar resistance under field conditions (Knüfer et al. 2016).

Emerging diseases pose threats to natural and cultivated ecosystems (Fisher et al. 2012). Pathogen emergence can be preceded by recent introductions if disease propagules reach unaffected regions. For example, rusts are especially notorious for their natural ability of long-distance dispersal as their wide-ranging airborne spread of spores makes that local rust populations can change swiftly. Hence, dramatic shifts in the European wheat yellow rust (*Puccinia striiformis* f. sp. *tritici*) populations have been observed in recent years as exotic lineages have replaced former populations (Hubbard et al. 2015; Hovmøller et al. 2015). Furthermore, disease dispersal is also facilitated through human activities. Globalization has led to more inter-continental trade and transport of humans and commodities, which, unquestionably, has contributed to an increasing frequency of pathogen introductions. For example, the introduction of the North American forest pathogen *Heterobasidion irregulare* into Italy coincided with an American military base in World War II (Garbelotto et al. 2013). However, recent introductions are not prerequisites for new disease emergences (Anderson et al. 2004). Environmental factors, such as weather and farming techniques, can give rise to established, yet latent, pathogen populations. Verticillium stem striping occurred in the United Kingdom (UK) in this fashion as a genetic diverse *V. longisporum* population suddenly emerged for currently unknown reasons (Depotter et al. 2017).

Verticillium stem striping was reported in 2007 for the first time in the UK and is currently widespread in England (Gladders et al. 2011, 2013). The recent and sudden rise of this disease makes that hitherto little is known about the pathogenic traits of the British *V. longisporum* population. The aim of this study is to assess the threat of Verticillium stem striping in the UK by comparing the pathogenicity British isolates with previously characterized *V. longisporum* strains from different countries.

## Materials and methods

### Pathogenicity tests

Pathogenicity tests were conducted in order to compare the virulence of 9 *V. longisporum* strains (Table 1). Three different *Brassica* crops were used: oilseed rape (*B. napus* var. *oleifera*), cauliflower (*B. oleracea* var. *botrytis*) and Chinese cabbage (*B. rapa* subsp. *Pekinensis*). Winter oilseed rape cultivars comprise open pollinating and hybrid types of which one of each was tested: cv. Quartz and cv. Incentive, respectively. Furthermore, one Chinese cabbage cultivar (cv. Hilton), and one cauliflower cultivar (cv. Clapton) were also included. Thus, in total, four *Brassica* cultivars were tested. Plants were grown in climate-controlled glasshouses under a 16 hrs light/ 8 hrs dark cycle with temperatures maintained between 20 and 28°C during the day, and a minimum of 15°C overnight. Before sowing, seeds were surface-sterilized during 1 min in 70% ethanol, followed by 15 min in 1% commercial bleach and then rinsed four times with sterile water. These sterilized seeds were then sown in trays with sterile compost and kept in the glasshouse for 14 days. These two-week old seedlings were inoculated by dipping the roots for 30 min in a suspension of 1×10^6^ conidiospores ml^−1^. Conidiospores were obtained from three-week old cultures grown on potato dextrose agar plates. Fifteen plants were inoculated for every *V. longisporum* strain and 15 control plants were dipped in sterile water instead of a conidiospore suspension. Individual seedlings were planted in 9 cm square pots with 4:1:1 compost:sand:loam mixture and pots were placed according to a random block design. The potting mixture had been autoclaved twice for 60 min with 24h between each treatment. Plants were grown for 6 weeks before harvesting. Plants were harvested by cutting the stem just above the hypocotyls and pooled in 5 groups of three plants. Above ground dry weight was subsequently determined by drying the samples at 100°C for a minimum of 12 h. Pathogenicity tests were executed twice to confirm the virulence responses of the different strains. In addition, stem samples were taken and pooled into 5 groups to determine the relative fungal colonization of the *V. longisporum* strains. Stem pieces of 1 cm, just above the hypocotyls, were cut. During the repeat experiment, Quartz plants were only grown until 25 days after inoculation to guarantee sufficient plant biomass to extract DNA from as longer growth periods could lead to complete decay of the plants (Figure S1). Furthermore, in the second pathogenicity test, Chinese cabbage plants were inoculated with a water suspension of 5×10^5^ conidiospores ml^−1^, instead of 1×10^6^ conidiospores ml^−1^.

**Table 1:**
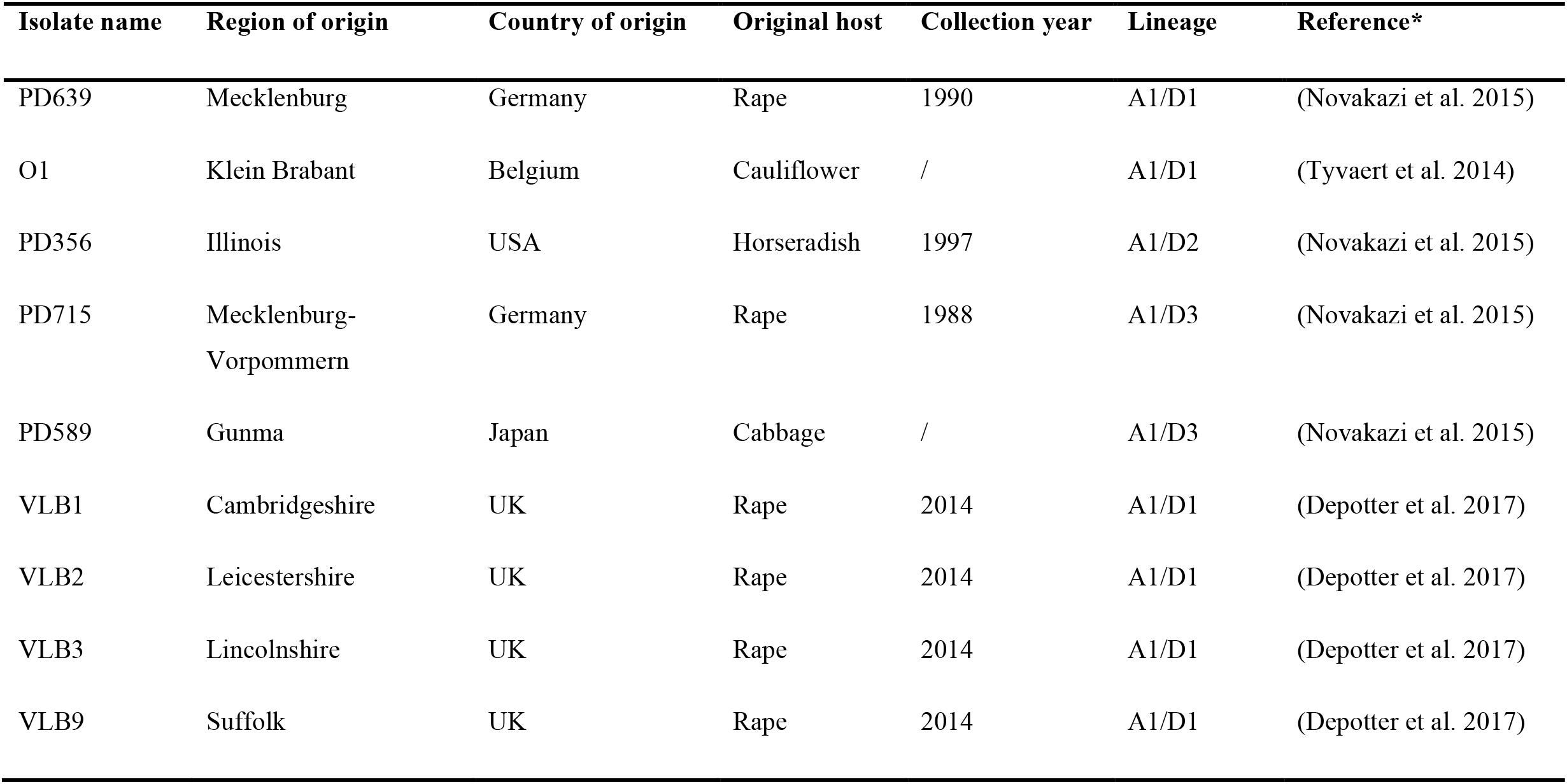
*Verticillium longisporum* isolate information.

| Isolate name | Region of origin | Country of origin | Original host | Collection year | Lineage | Reference* |
| --- | --- | --- | --- | --- | --- | --- |
| PD639 | Mecklenburg | Germany | Rape | 1990 | A1/D1 | (Novakazi et al. 2015) |
| O1 | Klein Brabant | Belgium | Cauliflower | / | A1/D1 | (Tyvaert et al. 2014) |
| PD356 | Illinois | USA | Horseradish | 1997 | A1/D2 | (Novakazi et al. 2015) |
| PD715 | Mecklenburg-Vorpommern | Germany | Rape | 1988 | A1/D3 | (Novakazi et al. 2015) |
| PD589 | Gunma | Japan | Cabbage | / | A1/D3 | (Novakazi et al. 2015) |
| VLB1 | Cambridgeshire | UK | Rape | 2014 | A1/D1 | (Depotter et al. 2017) |
| VLB2 | Leicestershire | UK | Rape | 2014 | A1/D1 | (Depotter et al. 2017) |
| VLB3 | Lincolnshire | UK | Rape | 2014 | A1/D1 | (Depotter et al. 2017) |
| VLB9 | Suffolk | UK | Rape | 2014 | A1/D1 | (Depotter et al. 2017) |

*Arabidopsis thaliana* (Col-0) plants were grown in a different glasshouse where the temperature was kept between 19 and 21°C and the same light/dark regime was used (16h/8h). Four plants of every treatment were grown. Plant inoculation occurred three weeks after sowing and roots were dipped for 10 min in a water suspension of 10^6^ conidiospores ml^−1^. Above-ground plant material was harvested for real-time PCR analysis three weeks after inoculation.

### Relative fungal quantification

Samples were taken for fungal biomass quantification and ground in liquid nitrogen. Approximately 200 mg ground plant material was dissolved in 500 μl of CTAB buffer (55 mM CTAB, 0.1 M Tris pH 8.0, 20 mM EDTA pH 8.0, 1.25 NaCl and 0.25 mM PVP 40). The buffer/plant extract was incubated for 30 min at 65°C. Samples were centrifuged and supernatant was subsequently transferred to a clean tube and 250 μl chloroform:isoamyl alcohol solution (24:1) was added. Samples were mixed thoroughly by inversion and subsequently centrifuged. Next, the supernatant was transferred into 50 μl ammonium acetate (7.5M) and 500 μl ethanol. After mixing, DNA was precipitated and the supernatant was removed. DNA pellets were washed twice with 70% ethanol and dissolved in DNase free water.

The amount of *V. longisporum* DNA in stems was quantified relatively to the amount of plant DNA by real-time PCR using QuantStudio™ Flex Real-Time PCR System (Applied Biosystems, CA, USA). Fungal DNA was amplified with the *V. longisporum* specific primer pair: n VlTubF2/ VlTubR1 (GCAAAACCCTACCGGGTTATG / AGATATCCATCGGACTGTTCGTA) (Debode et al. 2011) and the RuBisCO sequence targeting primer pair RubF/RubR (TATGCCTGCTTTGACCGAGA / AGCTACTCGGTTAGCTACGG) for plants. Real-time PCR was performed in reactions of 10 μl containing 500 nM of every primer and 5 μl of Power SYBR Green Master Mix (Applied Biosystems, Foster City, CA, USA). The thermal programme of the real-time PCR started with an initial denaturation step at 95°C for 10 min, followed by 40 cycles of 15 s at 95°C, 1 min at 62°C and 30 s at 72°C. Specific amplification was verified running a melting curve: samples were heated to 95°C for 15 s, cooled down to 60°C for 1 min and heated again to 95°C for 15 s. Signals above 36 cycles are considered below the detection limit.

DNA of the *Arabidopsis* plants was isolated according to Fulton et al. (1995). Relative *V. longisporum* colonization was quantified according to Ellendorf et al. (2009) using the qPCR core kit (Eurogentec, Liège, Belgium). Real-time PCR was executed on an ABI7300 PCR System (Applied Biosystems, Foster City, CA, USA) with following thermal conditions: an initial 95°C denaturation step for 4 min, 30 cycles of denaturation for 15 s at 95°C, annealing for 30 s at 60°C, and extension for 30 s at 72°C.

### Data analysis

Significance levels for parametric data were calculated with the one-way analysis of variance (ANOVA) or the t-test for unequal variances in R3.2.3 (R Core Team 2015). Non-parametric data was analysed with the Mann-Whitney U-test. Correlations were calculated with the Pearson correlation coefficient (*r*).

## Results

The pathogenicity of four British *V. longisporum* isolates was compared with that of five previously characterised isolates, including isolates of all three *V. longisporum* hybridization lineages (Table 1). The isolates were tested on four different *Brassica* cultivars: two oilseed rape (cv. Incentive and cv. Quartz), one cauliflower (cv. Clapton) and one Chinese cabbage (cv. Hilton). Pathogenicity tests were repeated twice and similar pathogenicity outcomes for the different *V. longisporum* strains were obtained on both occasions. Pathogenic *V. longisporum* isolates stunted plant growth and leaves displayed chlorosis and necrosis (Figure S1-4). Moreover, the more aggressive isolates caused complete decay of Quartz plants within 6 weeks of inoculation (Figure S1). Internally, vascular discoloration of diseased plant roots was observed, ranging from light brown to black. Furthermore, the pathogenicity of the British strains was also tested on the model plant *Arabidopsis thaliana* (Col-0): VLB1, VLB2 and VLB9 induced leaf curling and necrosis (Figure S5). The same symptoms were observed for VLB3, but the plants were also heavily stunted indicating that VLB3 is more aggressive than the other British strains on *Arabidopsis*.

The British *V. longisporum* isolates were pathogenic on all tested *Brassica* cultivars (Figure 1). Disease symptoms such as stunted growth and necrosis led to significant aboveground biomass reductions, which were amongst the highest for the *V. longisporum* strains tested. Similarly, severe biomass reductions were observed for the two A1/D1 representatives (PD639 and VLO1), illustrating the high virulence of this lineage on *Brassica* hosts (Figure 1). In contrast, PD356 was the least virulent *V. longisporum* strain, especially on oilseed rape, where no significant reductions in biomass accumulation were observed upon inoculation (Figure 1A-B). PD356 inoculation significantly reduced the plant biomass of Clapton and Hilton but nevertheless it was one of the least aggressive isolates (Figure 1C-D). Disease responses of the two A1/D3 representatives (PD715 and PD589) were, in contrast to the two A1/D1 representatives, dissimilar (Figure 1). The German isolate PD715 was a weak pathogen unable to cause significant biomass reduction on oilseed rape (Figure1 A-B) and was amongst the weakest strains tested on Clapton and Hilton (Figure 1C-D). In contrast, the Japanese A1/D3 isolate, PD589, strongly affected *Brassica* crops as PD589 was amongst the most severe isolates in Quartz, Clapton and Hilton (Figure 1A-C-D). Intriguingly, although the devastating outcome on the oilseed cultivars Quartz, no significant biomass reduction was observed when Incentive was infected with PD589 (Figure 1B).

**Figure 1:**
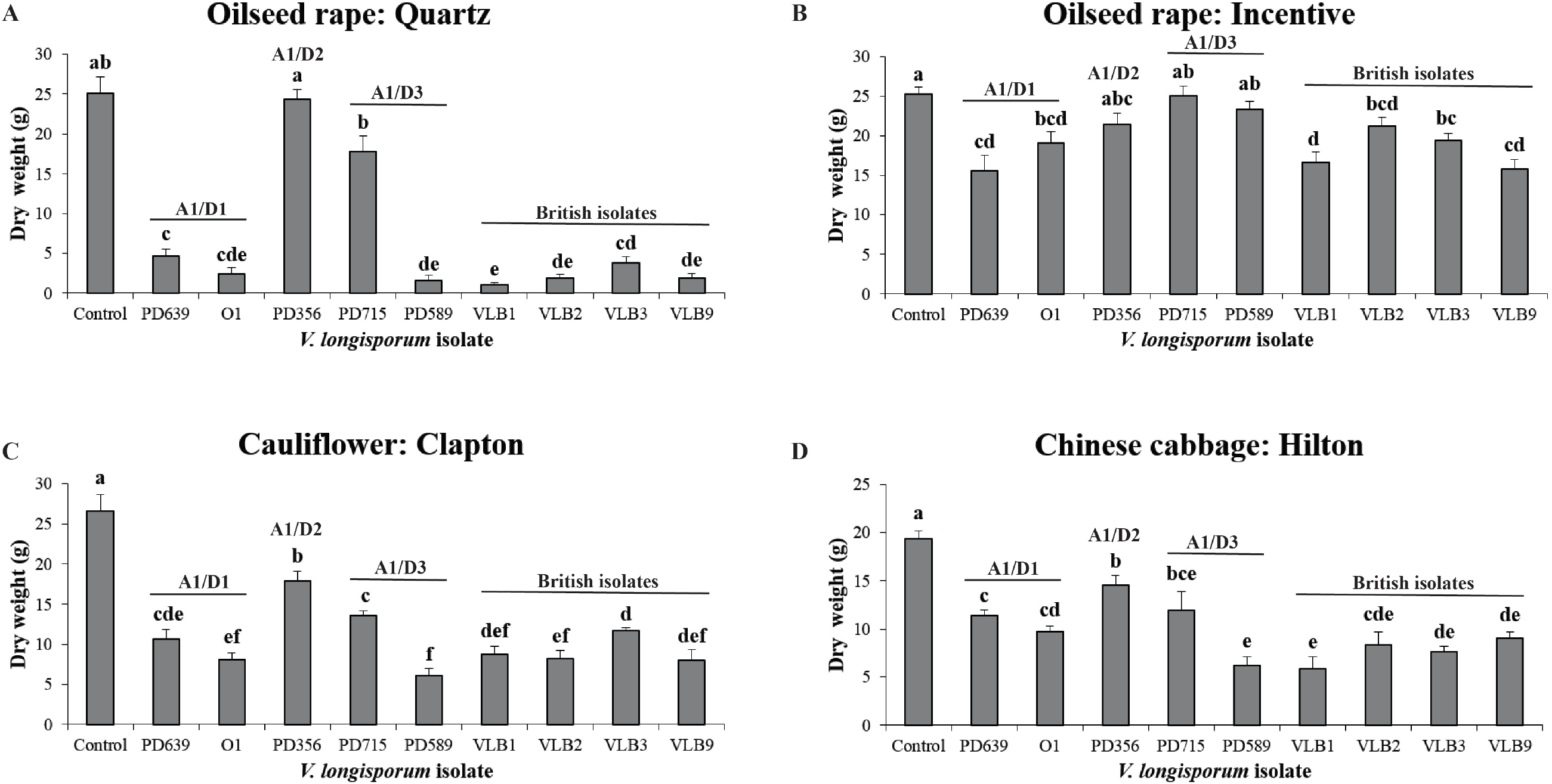
Aerial plant biomass accumulation of various Brassicaceae crops upon inoculation with *Verticillium longisporum*. The bars indicate the average above ground dry weight at 6 weeks after inoculation for the cultivars Quartz, Clapton and Hilton (A, C and D), whereas the bars for Incentive give the median weight (B). Significant levels were calculated for Quartz, Clapton and Hilton with the t-test for unequal variances (*p* < 0.05), whereas significance levels for Incentive were calculated Mann-Whitney U-test (*p* < 0.05). Error bars represent the standard error. Different letter labels indicate significant differences.

To determine to what extent the symptomatology correlates with the amount of *V. longisporum* biomass inside the plant, *V. longisporum* DNA was quantified inside the stems relative to the amount of plant DNA. Verticillium spp. are xylem colonizers, hence stem colonization is a good indication for strain aggressiveness. In correspondence with the observed disease symptoms, PD356 and PD715 were weak *Brassica* colonizers, as both strains were generally not detected in the stem, except for Hilton where PD715 was detected in four of the five samples (Figure 2). In contrast, all other *V. longisporum* isolates could be detected in most cases. In Quartz, PD589 was clearly the best colonizer, whereas the British and the A1/D1 strains were approximately present in equal levels (Figure 2A). The colonization of the detected isolates was negatively correlated with the aboveground biomass of Quartz plants (*r* = −0.6526, *p* = 2.148×10^−5^). In Incentive, significant differences in the colonization of the pathogenic isolates were observed, although these were not significantly correlated to plant biomass (*r* = −0.2745, *p* = 0.1105) (Figure 2B). Intriguingly, although no significant biomass reduction was observed upon infection of PD589 on Incentive (Figure 1B), PD589 was able to colonize the stem to a similar extent as the A1/D1 strains. In Clapton, high differences in colonization of the same isolates were observed between biological replicates (Figure 2C). In correspondence with Quartz and the disease symptoms, PD589 had the highest median colonization level, which was significantly higher than that of PD639 and VLB3. The fungal colonization of the detected isolates was also negatively correlated to the aboveground biomass of the cauliflower plants (*r* = −0.4793 *p* = 4,127 × 10^−3^). Hilton was the only *Brassica* cultivars with detection for PD715, however, no differences in colonization between the isolates were observed (Figure 2D). Similarly, PD715 was also detected in *Arabidopsis* for half of the inoculated plants and no significant differences in colonization were found between the treatments (Figure 3).

**Figure 2:**
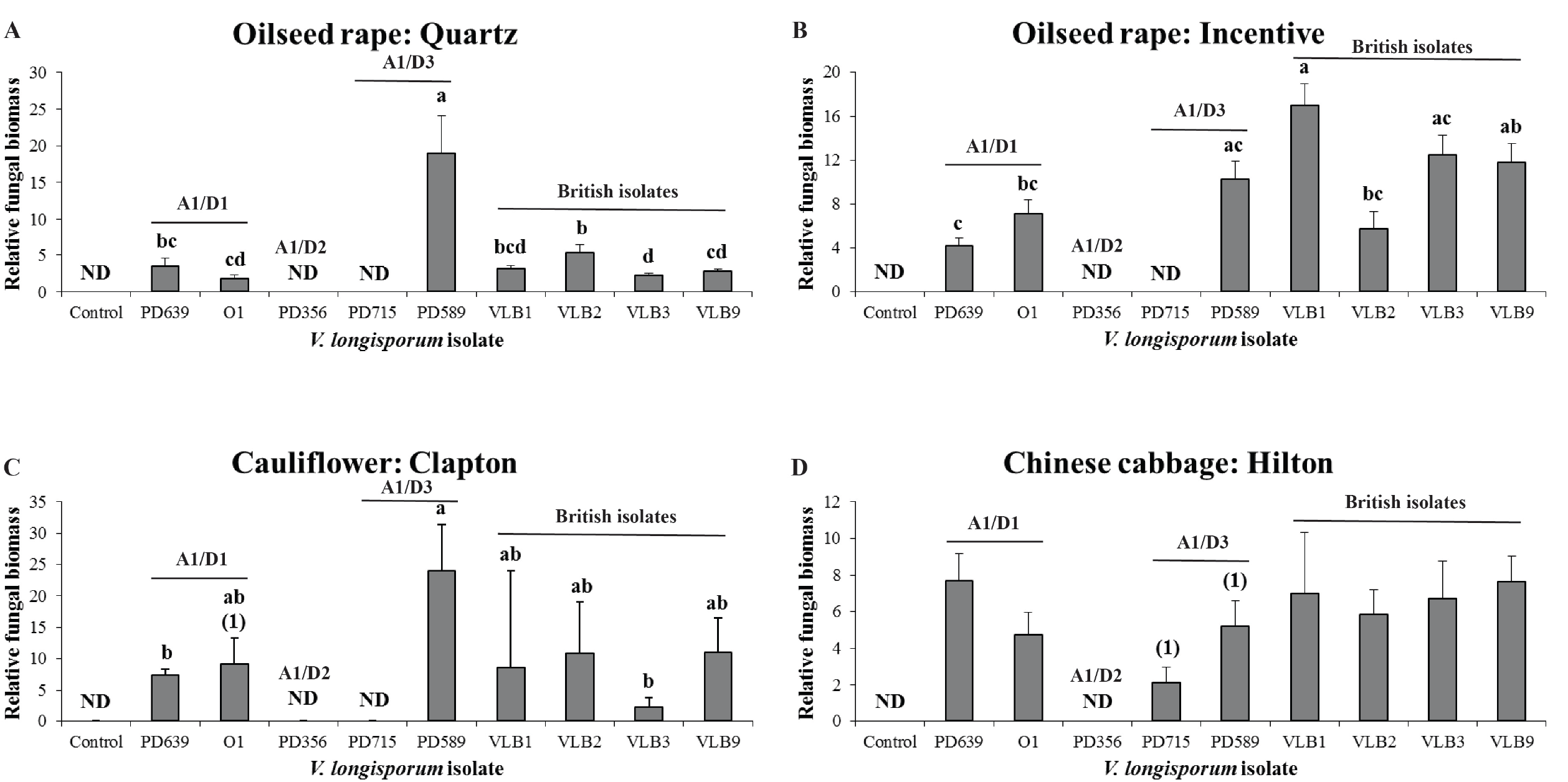
Fungal biomass accumulation of various *Verticillium longisporum* strains in plant stems. The bars indicate the median *V. longisporum* biomass relatively to the stem biomass. Isolates with no bar and ND in the graph were not detected in all 5 biological replicates. Significant differences were calculated with the Mann-Whitney U-test (*p* < 0.05) and depicted by different letter labels. No significant differences in colonization between isolates were found in Chinese cabbage cultivar Hilton. The number between brackets gives the amount of samples without detection. No number is given if the fungal colonization in all replicates was detected. Error flags represent the standard error.

**Figure 3:**
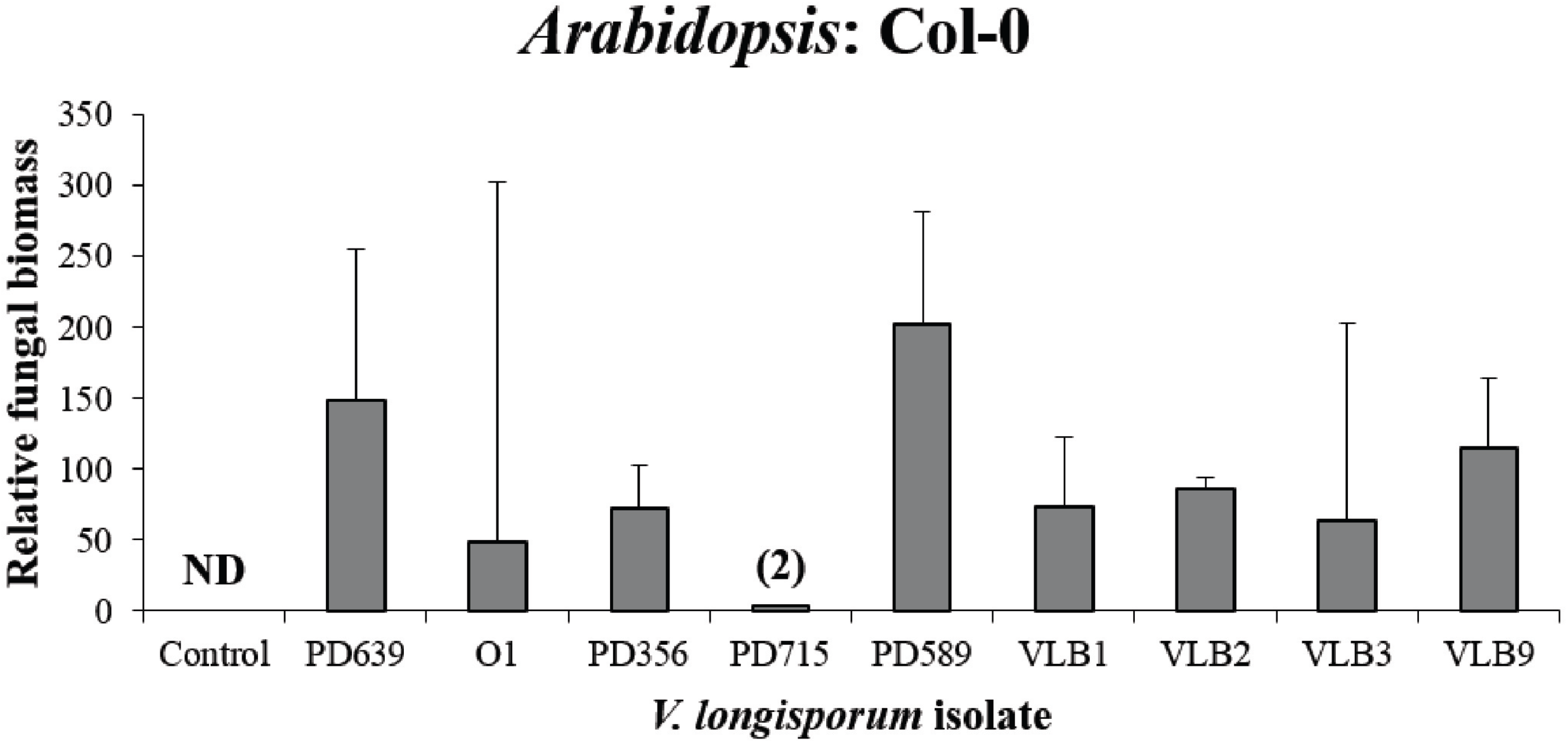
Fungal biomass accumulation of various *Verticillium* longisporum strains in *Arabidopsis* plants. The bars indicate the median *V. longisporum* biomass relatively to plant biomass. Isolates with no bar and ND in the graph were not detected in all 5 biological replicates. No significant differences in colonization of the different isolates were found (Mann-Whitney U-test, *p* < 0.05). The number between brackets gives the amount of samples without detection. No number is given if the fungal colonization in all replicates was detected. Error flags represent the standard error.

## Discussion

Emerging diseases are often received with concern as their rise is threatening and their final impact uncertain. Hence, recent emergences of the humane virus diseases Ebola in 2014 and Zika in 2015 led to global unsettlement (Attar 2016; Meyers et al. 2015). The majority of emerging diseases originate from the introduction of pathogens in new geographic regions (Anderson et al. 2004). Here, comparisons between emerging pathogens and their original source can help to indicate to what extent the emerging disease may impact ecosystems. However, environmental factors and ecosystem management can lead to sheer differences in the disease outcome of the same pathogen. The recent outbreak of Verticillium stem striping in the UK exemplifies the far-reaching consequences of environmental and anthropogenic factors as the disease originates from a latent, established *V. longisporum* population (Depotter et al. 2017). Changes in weather conditions and/or oilseed rape cultivation are likely to have favoured conditions that supported the emergence of a British *V. longisporum* population capable of initiating new disease epidemics. Similar to other newly emerging diseases, Verticillium stem striping it threatening for British oilseed rape, as the impact on yield is still relatively uncertain. In this study the pathogenicity of four British *V. longisporum* isolates from different counties was compared with five previously characterized strains including strains from all *V. longisporum* lineages (Table 1). The British isolates caused significant reduction in the aerial biomass of all the cultivars tested and colonized plants successfully (Figure 1–3). The disease level of British isolates resembled those of the other two A1/D1 lineage isolates (PD639 and VLO1) as plant colonization caused significant yield reductions on all *Brassica* crops (Figure 1). *V. longisporum* lineage A1/D1 can therefore generally be considered pathogenic on *Brassica* crops. This corresponds with a previous *V. longisporum* pathogenicity test that considered A1/D1 as the most virulent *V. longisporum* lineage on oilseed rape and cauliflower (Novakazi et al. 2015).

Lineage A1/D2 has hitherto only been found on horseradish and causes the severest disease on this crop of all *V. longisporum* strains (Novakazi et al. 2015). Isolate PD356 is a relatively poor colonizer of *Brassica* crops and caused no or only a small reduction in plant biomass (Figure 1–2). The strong disease symptoms on horseradish and the lack of disease in *Brassica* crops indicated that the host range of lineage A1/D2 is rather specialized. However, A1/D2 came out as the most virulent *V. longisporum* lineage on cabbage (B. *oleracea* convar. *capitata* var. *alba,* cv. Brunswijker) during pathogenicity tests of Novakazi *et al.* (2015) indicating that pathogenicity may vary within different varieties or crop sub-species. Thus, generalizations about the pathogenicity of the *V. longisporum* lineages should be treated with caution. Lineage A1/D3 was considered non-pathogenic on oilseed rape (Zeise and Tiedemann 2002). However, this pathogenicity study showed, along with a previous study, that non-pathogenicity of lineage A1/D3 cannot be generalized on oilseed rape (Novakazi et al. 2015). Lineage A1/D3 isolate PD715 was non-pathogenic on oilseed rape and was also not detected in the stem after inoculation. In contrast, PD589 is a strong *Brassica* stem colonizer (Figure 2) and caused severe disease symptoms in Quartz (Figure 1A). Intriguingly, this was in contrast with the symptoms on Incentive that, despite extensive stem colonization, tolerated infection by PD589.

Interspecific hybridization is an intrusive evolutionary mechanism that alters host range and virulence of pathogens (Depotter et al. 2016b). Hybrid pathogens can therefore have devastating outcomes on ecosystems. For example, the hybrid pathogen *Phytophthora xalni,* lead to an epidemic of alder decline in riparian ecosystems of Europe since the early 1990s (Brasier et al. 1995). The hybrid fungus *V. longisporum* is currently gaining momentum as it is causing Verticillium stem striping into previously unaffected regions (Gladders et al. 2011; CFIA 2015). A decade since the first report of Verticillium stem striping in the UK, the extent to which this disease may contribute to losses in British oilseed rape is better understood; British *V. longisporum* strains are as pathogenic as other A1/D1 strains isolated from various geographical regions and given similar climatic constraints at these locations, Verticillium stem striping is therefore expected to have similar outcomes as in countries where the disease was previously established such as Germany and Sweden. However, the British *V. longisporum* population is genotypically more diverse than the ones in Germany and Swedish (Depotter et al. 2017). Conceivably, the heterogeneous character of the British populations may therefore hamper disease management strategies more. Nevertheless, these recent findings should be an incentive in oilseed rape breeding programs to select for Verticillium stem striping resistance, especially as protective or curative control by conventional fungicides is not possible for *Verticillium* diseases.

## Acknowledgments

The authors would like to thank the Marie Curie Actions programme of the European Commission that financially supports the research investigating the threat of *V. longisporum* to UK oilseed rape production. Work in the laboratory of B.P.H.J.T. is supported by the Research Council Earth and Life Sciences (ALW) of the Netherlands Organization of Scientific Research (NWO).

